# Complete Telomere-to-Telomere Assembly of the Y Chromosome in the Chinese Quartet

**DOI:** 10.64898/2026.04.13.718326

**Authors:** Bo Wang, Zexin Zhang, Shijie Wan, Pengyu Zhang, Yu Zhang, Xia Wang, Lianhua Dong, Kai Ye, Xiaofei Yang

**Affiliations:** School of Automation Science and Engineering, Faculty of Electronic and Information Engineering, Xi’an Jiaotong University, Xi’an 710049, China; MOE Key Lab for Intelligent Networks & Networks Security, Faculty of Electronic and Information Engineering, Xi’an Jiaotong University, Xi’an 710049, China; School of Computer Science and Technology, Faculty of Electronic and Information Engineering, Xi’an Jiaotong University, Xi’an 710049, China; Center for Advanced Measurement of Science, National Institute of Metrology, Beijing 100029, China; Center for Reference materials Research and Management, National Institute of Metrology, Shenzhen 518107, China; Faculty of Science, Leiden University, Leiden 2311 EZ, The Netherlands

## Abstract

The complete assembly of the human Y chromosome remains a formidable task due to its highly repetitive, palindromic, and heterochromatic architecture. While complete telomere-to-telomere (T2T) assemblies have recently been generated for select individuals, high-quality, gapless Y chromosome references for East Asian lineages—specifically within well-characterized multi-omics reference cohorts—remain scarce. The Chinese Quartet (CQ), comprising monozygotic twin daughters and their parents, represents a premier national reference material system for genomic standardization. However, because the female offspring inherited their father’s X chromosome rather than his Y chromosome, previous pedigree-based T2T assemblies (CQ v3.2) lacked a gapless Y chromosome sequence. Here, we present a complete, gapless T2T assembly of the Y chromosome (designated CQ_chrY) from the father of the CQ. This assembly was generated by integrating Oxford Nanopore Technologies ultra-long reads, PacBio high-fidelity reads, and Hi-C data, resulting in a sequence of 68.50 Mb. The assembly shows exceptional base accuracy (QV = 57.54) and structural assembly quality (CRAQ AQI = 99.781). We completely resolved the 40.09 Mb Yq12 heterochromatic region, a ∼940 kb centromeric higher-order repeat array, and annotated 96 protein-coding gene models evaluated against GENCODE/T2T-Y standards, alongside 57.59 Mb (84.07%) of repetitive elements. CQ_chrY serves as a complementary paternal resource extending the CQ reference platform, providing a robust foundation for population genetics, structural variation analysis, and East Asian genomic standardization.

## Background & Summary

The human Y chromosome plays an essential role in male sex determination, spermatogenesis, and fertility, yet it has long represented the most structurally challenging and incompletely assembled portion of the human genome. Approximately 95% of its length lies within the non-recombining male-specific region, which is densely packed with complex repetitive architectures, including segmental duplications, ampliconic palindromes, satellite arrays, and massive heterochromatic blocks^1^. Short-read sequencing technologies were inherently incapable of spanning these extensive repetitive domains, leaving over half of the Y chromosome sequence unresolved as sequence gaps in standard reference assemblies such as GRCh38^2^. Consequently, crucial research into paternal genetic variation, male-specific disease susceptibility, and human evolutionary lineage history was substantially constrained.

The emergence of long-read sequencing technologies, specifically PacBio high-fidelity (HiFi) reads and Oxford Nanopore Technologies (ONT) ultra-long (UL) reads, has fundamentally transformed *de novo* genome reconstruction^3,4^. In 2022, the Telomere-to-Telomere (T2T) Consortium achieved the first gapless assembly of a human complete genome (CHM13), though it lacked a Y chromosome^5^. This achievement was extended in 2023 by the complete T2T assembly of the HG002 Y chromosome, which resolved the remaining approximately 30 Mb of heterochromatic genome regions and established a baseline for Y chromosome structural biology^6^. Simultaneously, the Human Genome Structural Variation Consortium (HGSVC) expanded this landscape by assembling 43 diverse Y chromosomes^7^, revealing that the Y chromosome is among the most rapidly evolving human chromosomes, exhibiting dramatic length variations predominantly driven by copy-number fluctuations in the Yq12 heterochromatic block and centromeric satellite arrays. These insights underline that a single reference sequence cannot capture the extensive structural diversity present across diverse human lineages.

The Chinese Quartet (CQ) family—comprising a father, mother, and monozygotic twin daughters—serves as a primary multi-omics reference material system (GBW09900–GBW09903, authorized by the State Administration for Market Regulation, China) widely adopted for benchmarking genomic tools and variant calling pipelines^8,9^. In previous pedigree-based T2T assembly efforts (CQ v3.2), maternal and paternal haplotypes were resolved using pooled reads from the twin daughters^10^. Because the female offspring inherited their father’s X chromosome rather than his Y chromosome, the paternal Y chromosome was inherently absent from the CQ v3.2 diploid reference^10^. To address this limitation, the current study presents a complete, *de novo* T2T assembly of the paternal Y chromosome (CQ_chrY) generated directly from the father’s own cell line (LCL7, GBW09900).

CQ_chrY v1.3 spans 68,501,441 bp from the 5’ pseudoautosomal region 1 (PAR1) telomere to the 3’ PAR2/q-terminal telomere without a single internal gap. The assembly exhibits a quality value (QV) of 57.54, and a Clipping Reveals Assembly Quality (CRAQ) overall Assembly Quality Indicator (AQI) of 99.781. Key structural milestones include the complete resolution of the 40.09 Mb Yq12 heterochromatic block and the precise characterization of a ∼940 kb centromeric higher-order repeat array. As the third complete Chinese Y chromosome assembly reported to date (alongside CN1 and YAO)^11,12^, CQ_chrY complements the existing CQ v3.2 reference system and provides a high-quality East Asian reference for comparative genomics and paternal lineage evolution^13^.

## Methods

### Sample collection and cell line characterization

The sample evaluated in this study originated from a B-lymphoblastoid cell line (LCL7, national reference material accession GBW09900) established from the 60-year-old father of the Chinese Quartet family. Cell line derivation, immortalization, and certification protocols were conducted under national reference material standards as detailed previously^14^. Liquid nitrogen-cryopreserved LCL7 cells were maintained at the National Institute of Metrology prior to high-molecular-weight (HMW) genomic DNA extraction.

### ONT ultra-long library preparation and sequencing

HMW genomic DNA was isolated using an SDS-based lysis protocol optimized to minimize physical shear forces. DNA integrity and concentration were validated via 0.75% agarose gel electrophoresis, NanoDrop 2000 spectrophotometry, and Qubit 3.0 Fluorometry (Invitrogen, Carlsbad, CA, USA). Crucially, no mechanical fragmentation or Covaris ultrasonic shearing was applied during library construction. HMW DNA fragments were size-selected using the BluePippin system (Sage Science, Beverly, MA, USA) with a high-pass cut-off.

End-repair and dA-tailing were performed using the NEBNext FFPE DNA Repair Mix and NEBNext Ultra II End Repair/dA-Tailing Module (New England Biolabs, Ipswich, MA, USA). Sequencing adapters were ligated using the SQK-LSK112 Kit (Oxford Nanopore Technologies, Oxford, UK). The resulting library was purified and loaded onto primed R10.4.1 flow cells and sequenced on a PromethION sequencer for 48 h at Xi’an Herui Gene Technology Co., Ltd. Base-calling was executed using Oxford Nanopore Dorado (v0.9.0, model dna_r10.4.1_e8.2_400bps_sup@v4.3.0) under super-accuracy (SUP) mode. Sequencing yielded 106.44 Gb of ultra-long read data, achieving a read N50 of 131.74 kb, a maximum read length of 1.33 Mb, and 66.06 Gb of yield in reads exceeding 100 kb (**Table 1**).

**Table 1.**
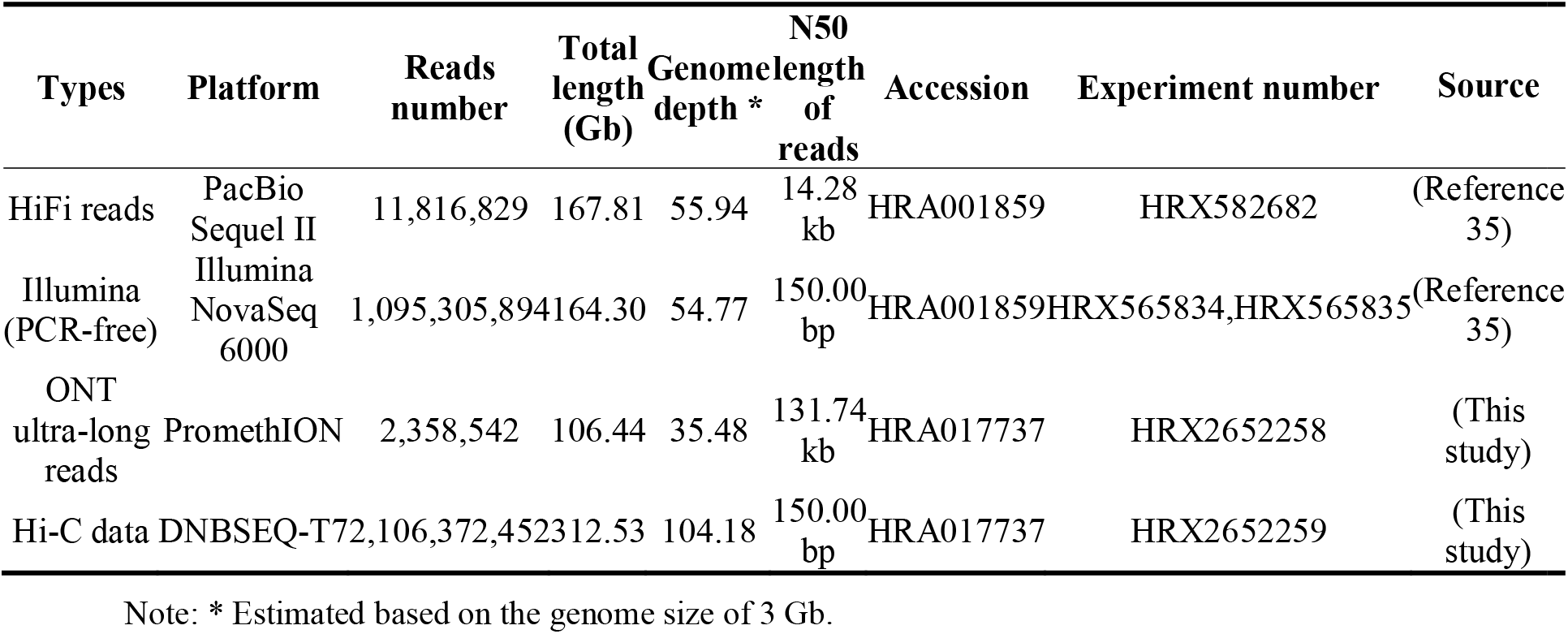
Summary of DNA sequencing data.

### Hi-C chromatin conformation capture sequencing

Hi-C libraries were prepared from cross-linked LCL7 cells following standard chromatin conformation capture procedures. Chromatin was digested with DpnII, biotin-labeled, and ligated. Library concentration and insert distribution were assessed using a Qubit 2.0 fluorometer and Agilent 2100 Bioanalyzer, with final quantification conducted via qPCR. Sequencing was performed on the DNBSEQ-T7 platform, generating 312.53 Gb of paired-end short reads across 1,053,186,226 total read pairs (**Table 1, Table S1**). Quality filtering demonstrated high library quality, with an alignment rate of 91.12% (959,698,345 read pairs) and a low PCR duplicate rate of 11.28% (**Table S1**). In total, 666,740,908 valid Hi-C contacts were identified using Juicer v1.6^15^ (63.31% of total read pairs and 79.29% of unique read pairs) (**Table S1**). For the assembled Y chromosome specifically, 302,5106 cis valid pairs were mapped (44,161 valid pairs per Mb), achieving an effective contact-map resolution of ≥ 10 kb (∼442 contacts per 10-kb bin; **Table S1**).

### *De novo* assembly workflow

The complete gapless T2T Y chromosome (CQ_chrY v1.3) was reconstructed using a fully *de novo*, multi-technology assembly and gap-filling strategy:

**1)** Initial whole-genome *de novo* assembly (**Figure S1**). All raw ONT UL reads were assembled *de novo* using hifiasm v0.25.0 in “--ont” mode^16^. This generated initial unitig graphs, yielding primary contigs covering chromosome Y. These primary unitigs were ordered and oriented against CHM13v2Y using RagTag v2.0.1^17^ to construct an initial draft scaffold (CQ_chrY v1.2) spanning ∼61.5 Mb with terminal telomeric repeat arrays identified at both ends (10,501 bp at the 5’ end and 1,817 bp at the 3’ end).
**2)** Gap-filling strategy and breakpoint validation (**Figure S2**). The initial draft assembly CQ_chrY v1.2 contained a single 100-bp central gap. Integration of PacBio HiFi, ONT UL reads, and Hi-C data assembled with hifiasm v0.25.0 generated a gap-resolving unitig, utg000737l (13.84 Mb). Using minimap2 v2.30^18^ (MAPQ = 60), utg000737l was uniquely anchored via a 3.62 Mb upstream anchor and a 10.19 Mb downstream anchor to replace the target gap-containing interval. Integrative

Genomics Viewer (IGV) tracks demonstrated seamless sequence continuity across both the upstream (pos: 27,564,000–27,566,000) and downstream (pos: 41,399,500–41,401,500) replacement breakpoints, validated by aligning HiFi and ONT UL reads with minimap2 v2.30 and Winnowmap v2.03^19,20^. The resulting CQ_chrY v1.3 assembly spans 68,501,441 bp and achieves gapless, T2T completeness (**Table 2**).

**Table 2.** Summary statistics of the Y chromosome.

| Item | Y chromosome |
| --- | --- |
| Size of assembly (Mb) | 68.50 |
| GC content (%) | 35.92 |
| Quality value | 57.54 |
| CRAQ AQI | 99.781 |
| CRAQ R-AQI | 100.00 |
| CRAQ S-AQI | 99.56 |
| Repetitive sequences (%) | 84.07 |
| Number of protein-coding genes | 96 |

### Regional annotation of the CQ_chrY Assembly

To comprehensively delineate the structural domains of CQ_chrY v1.3 (**Figure 1**), we adopted the standardized Y chromosome regional annotation pipeline developed by the T2T consortium (https://github.com/arangrhie/T2T-chrY). To ensure precise boundary definitions, pseudoautosomal regions (PAR1 and PAR2) underwent rigorous manual validation: the annotated PAR sequence was extracted and aligned against the chrX_RagTag assembly—generated by hifiasm v0.25.0 integrating ONT UL, PacBio HiFi, and Hi-C data—and manually inspected using Gepard v2.1^21^ dot plots. Ampliconic regions and palindromic units were similarly verified through manual inspection of local self-alignment dot plots. Furthermore, a dot-plot alignment between CQ_chrY v1.3 and CHM13v2Y confirmed high overall structural consistency between the two assemblies (**Figure S3**).

**Figure 1.**
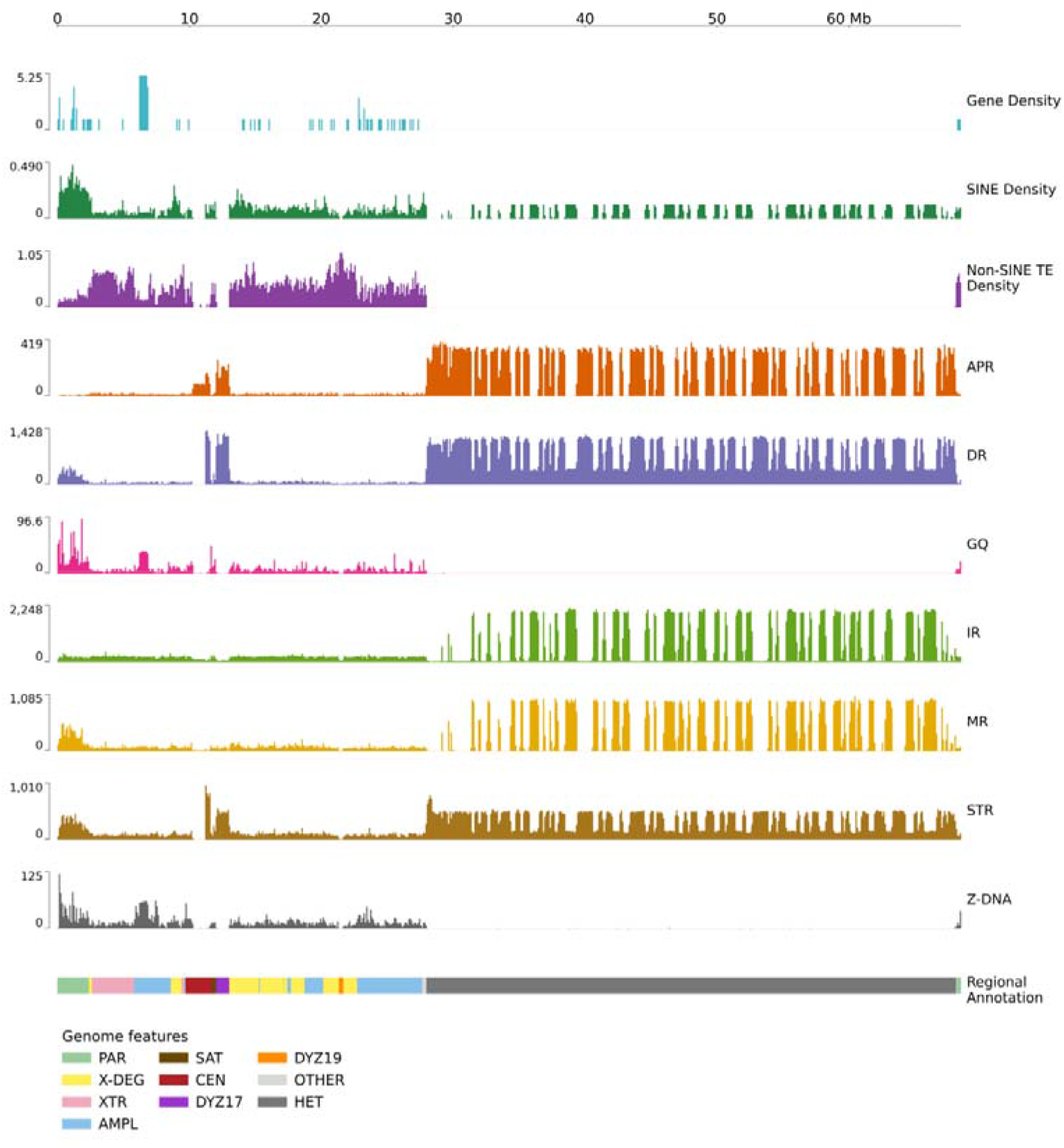
Structure of a complete CQ_chrY assembly. The gene density plot illustrates the distribution of protein-coding genes across the Y chromosome. SINE TEs are predominantly enriched in PAR1, whereas non-SINE TEs are largely restricted to the euchromatic region. Quantitative tracks display the distributions of various non-B DNA structures and repeat classes, including A-phased repeats (APR), direct repeats (DR), G-quadruplexes (GQ), inverted repeats (IR), mirror repeats (MR), short tandem repeats (STR), and Z-DNA. The bottom panel depicts regional annotations outlining major genomic features, such as pseudoautosomal regions (PAR), X-degenerate (X-DEG), X-transposed regions (XTR), ampliconic regions (AMPL), satellites/centromeres (SAT, CEN, DYZ17, DYZ19), and heterochromatin (HET).

### Pairwise collinearity and structural variation analysis

Pairwise collinearity between each Chinese Y assembly (CQ_chrY, CN1_chrY, and YAO_chrY) and the CHM13v2Y reference genome was evaluated using SyRI v1.7.0^22^. Chromosomes were first aligned with minimap2 to produce alignment SAM files, which were subsequently processed by SyRI v1.7.0 under default settings to identify syntenic regions, inversions, translocations, duplications, and non-aligned structural variations. Structural differences and pairwise chromosomal synteny across assemblies were visualized using plotsr v1.2.0^23^ (**Figure 2**). Pairwise whole-chromosome dot plots comparing CQ_chrY v1.3 (68,501,441 bp) against CN1_chrY v1.0.1 (65,662,189 bp) and YAO_chrY v2.0 (51,545,170 bp). Both comparisons display high sequence collinearity across the euchromatic region (**Figure S4**), interspersed with localized structural rearrangements along the main diagonal.

**Figure 2.**
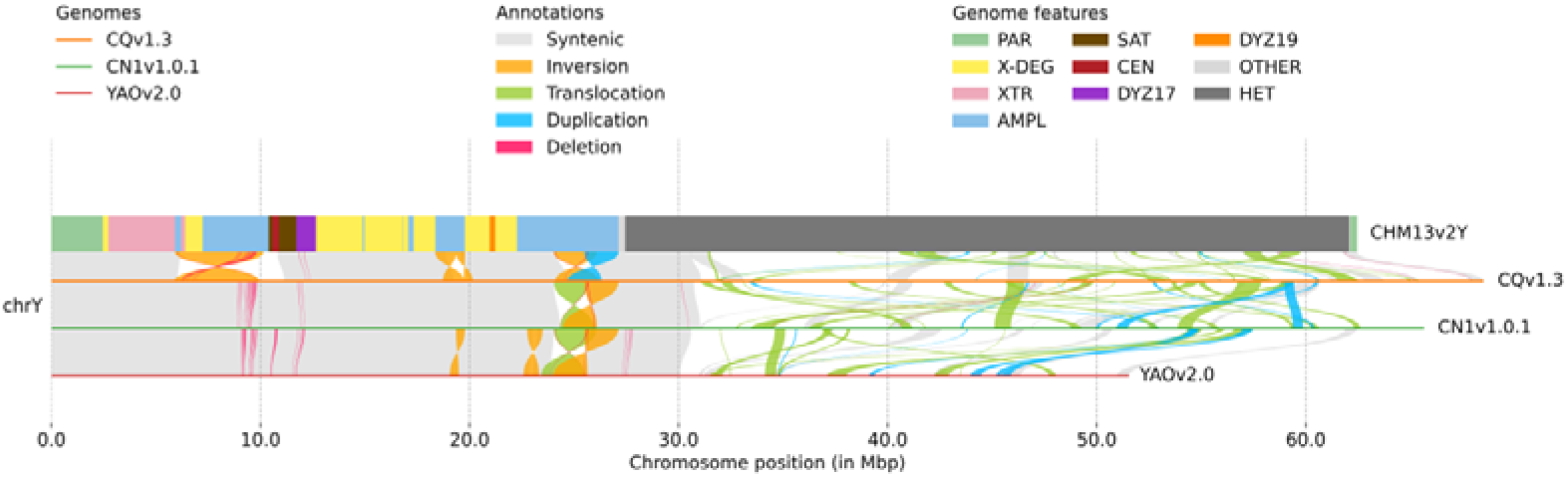
Comparative synteny and structural variations across Y chromosome assemblies. Genome alignment and structural variants among three Y chromosome assemblies (CQv1.3, CN1v1.0.1, and YAOv2.0) mapped against the CHM13v2Y reference genome. The top track displays the genomic features annotated along CHM13v2Y. Colored ribbons connecting the assemblies highlight distinct structural annotation types, including syntenic regions (light grey), inversions (orange), translocations (green), duplications (cyan), and deletions (pink). Chromosomal coordinates are shown along the horizontal axis in megabase pairs. PAR, pseudoautosomal region; X-DEG, X-degenerate region; XTR, X-transposed region; AMPL, ampliconic region; SAT, satellite; CEN, centromere; HET, heterochromatin.

The lower-right quadrant depicts the complex grid-like repeat patterns of the Yq12 heterochromatic block, demonstrating substantial length polymorphism and architectural variation among CQ, CN1, and YAO assemblies.

### Centromeric higher-order repeat architecture

CenMAP v0.2.4^24^, a centromere mapping and annotation pipeline, was employed to identify potential centromeric regions on the CQ_chrY genome. The α-satellite sequences were annotated using the script hmmer-run.sh (from https://github.com/fedorrik/HumAS-HMMER_for_AnVIL) with the HMM profile “AS-HORs-hmmer3.3.2-120124.hmm”. Subsequently, HOR structural variants (StV) were detected using the StV tool (https://github.com/fedorrik/stv). This analysis demonstrated that while the centromere is predominantly composed of 34-mer HOR units, CQ_chrY is notably characterized by the complete absence of the 36-mer HOR variant observed in CHM13v2Y, CN1, and YAO, reflecting lineage-specific centromeric divergence (**Figure 3**).

**Figure 3.**
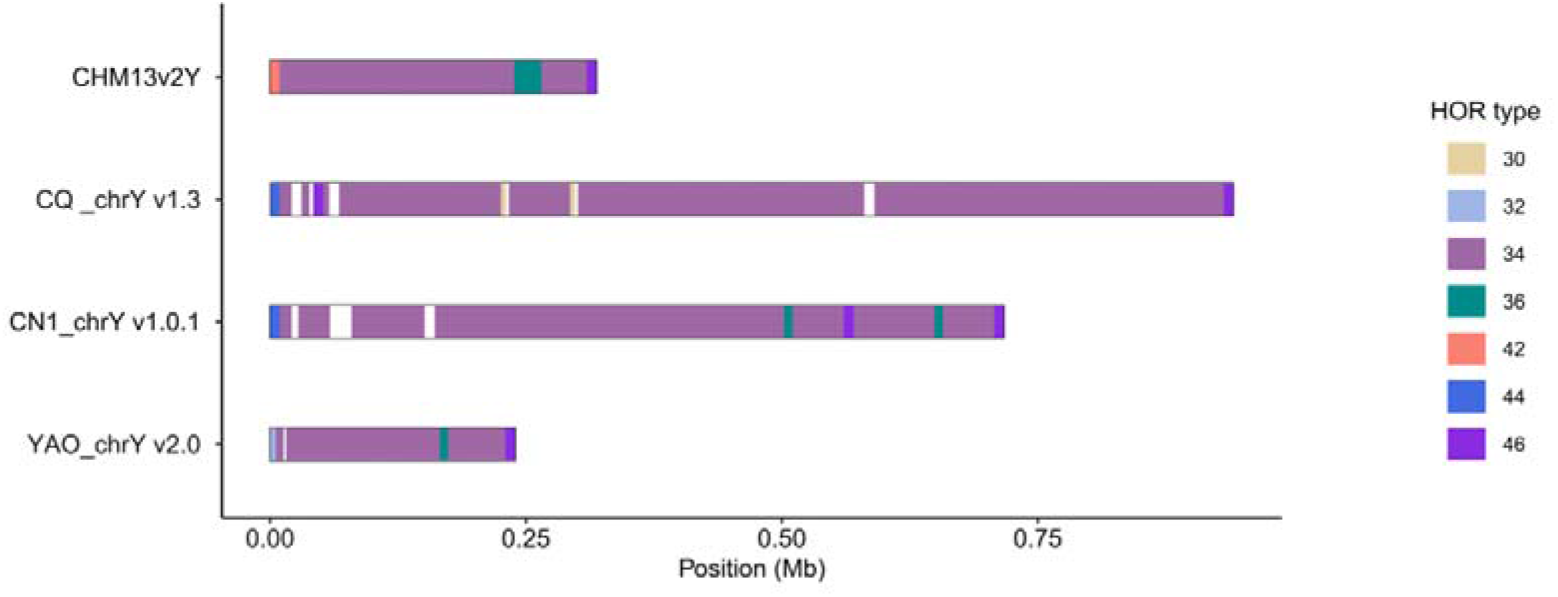
Comparative analysis of centromeric HOR organization across Y chromosome assemblies. The centromeric higher-order repeat (HOR) array varies significantly in length among assemblies, with CQ_chrY v1.3 spanning approximately 940 kb—nearly three times longer than CHM13v2Y (317 kb)—while CN1_chrY v1.0.1 and YAO_chrY v2.0 measure 715 kb and 238 kb, respectively. All four assemblies are predominantly composed of 34-mer HORs (purple). Notably, the 36-mer HOR variant (teal) is present in CHM13v2Y, CN1_chrY, and YAO_chrY, but completely absent in CQ_chrY v1.3. Other low-frequency HOR types (30-, 32-, 42-, 44-, and 46-mers) exhibit assembly-specific patterns along the centromeric regions.

### Composition and architecture of the Yq12 heterochromatic region

The Yq12 heterochromatic block was annotated using an HMM-based filtering pipeline (https://github.com/Markloftus/PanYChromosome) and completely resolved as a single, continuous 40,085,168 bp (∼40.09 Mb) sequence (**Figure 4, Table S2**). Architectural profiling revealed a conserved proximal boundary anchored by DYZ18 repeats, followed by extensive interspersed arrays of satellite repeats DYZ1 and DYZ2. CQ_chrY represents one of the largest gapless Yq12 heterochromatic blocks assembled in Chinese, spanning ∼40.09 Mb compared to 37.47 Mb in CN1 and 23.79 Mb in YAO.

**Figure 4.**
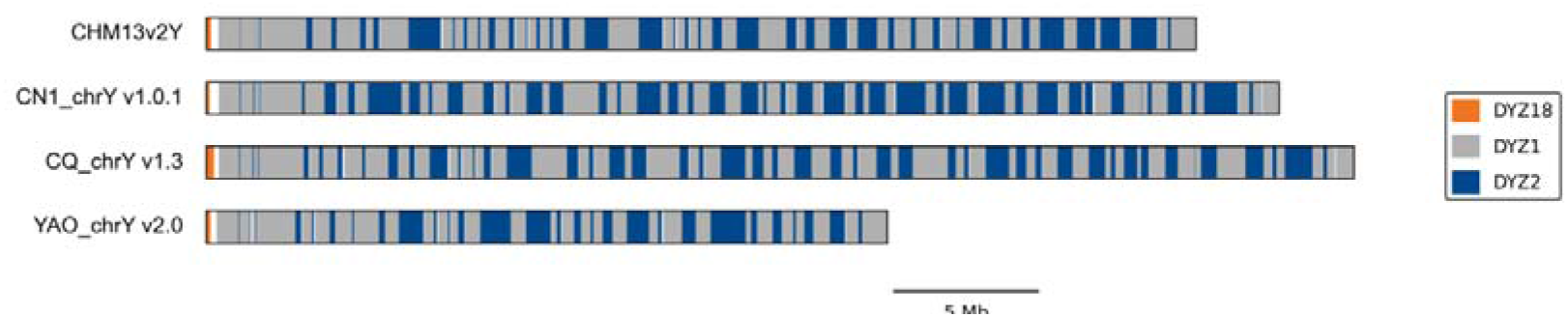
Composition and architecture of the Yq12 heterochromatic region across Y chromosome assemblies. Structural organization and repeat composition of the Yq12 heterochromatic region compared among CHM13v2Y, CN1_chrY v1.0.1, CQ_chrY v1.3, and YAO_chrY v2.0. All four assemblies display a conserved proximal boundary marked by DYZ18 (orange). The main body of the Yq12 block consists of interspersed arrays of satellite repeats DYZ1 (grey) and DYZ2 (dark blue). Substantial size polymorphism is observed across the assemblies, with CQ_chrY v1.3 possessing the longest Yq12 region and YAO_chrY v2.0 showing the shortest block. Scale bar, 5 Mb.

### Repetitive and Non-B sequence annotation

Repetitive sequences were annotated using RepeatMasker v4.1.7^25^ with the parameters “-e ncbi -species human”. The composition of repetitive sequences within CQ_chrY was comprehensively annotated (**Table 3**). The non-B DNA structures (G-quadruplexes, Z-DNA, A-phased repeats, direct repeats, mirrored repeats, and short tandem repeats) within the flanking sequences were identified using the nonB-GFA tool^26^. Non-B DNA track analysis revealed that regions with the potential to form alternative DNA secondary structures are highly enriched within satellite repeat arrays and the Yq12 heterochromatic region (**Figure 1**).

**Table 3.** Statistics of repetitive sequences in the Y chromosome assembly.

| Type | Length (bp) | % of genome |
| --- | --- | --- |
| SINEs | 4,599,866 | 6.71 |
| LINEs | 6,401,177 | 9.34 |
| LTR elements | 4,636,314 | 6.77 |
| DNA elements | 453,677 | 0.66 |
| Unclassified | 22,302 | 0.03 |
| Small RNA | 9424 | 0.01 |
| Satellites | 16,706,538 | 24.39 |
| Simple repeats | 24,669,219 | 36.01 |
| Low complexity | 101,957 | 0.15 |

### Gene annotation of CQ_chrY v1.3

To accurately map the gene landscape of CQ_chrY v1.3 while resolving complex multi-copy gene structures, gene annotation was conducted by implementing the T2T consortium-recommended pipeline^7^, leveraging Liftoff v1.6.3^27^ guided by the canonical CHM13v2Y reference annotation dataset (JHU RefSeq + Liftoff v5.3; https://github.com/marbl/CHM13). To prevent misannotation within highly duplicated ampliconic regions, parameters were set to allow copy number variation (-copies -sc 0.8). In total, 96 protein-coding genes were annotated across CQ_chrY v1.3 (**Table S3**). Future transcriptomic sequencing and functional experiments will be required to validate their expression profiles, pseudogenic status, and biological roles.

### Data Records

All raw sequencing datasets and the finalized gapless chromosome Y assembly (CQ_chrY v1.3) generated in this study have been deposited in public repositories under open access. Raw sequencing data are hosted in the Genome Sequence Archive for Human (GSA-Human)^28^ at the National Genomics Data Center (NGDC), China National Center for Bioinformation (CNCB), under accession number HRA017737^29^. The finalized T2T assembly sequence is deposited in GeneBase^30^ (NGDC) under accession number C_AA501122.1^31^.

The deposited datasets encompass ONT UL reads (85.56 GB) and paired-end Hi-C short reads (140.62 GB across two runs) in compressed FASTQ format (.fastq.gz / .fq.gz), as well as the complete genome assembly in compressed FASTA archive format (.fasta). The coordinate system for all downstream annotations and structural variant calls is defined by the gapless T2T *de novo* coordinate space of CQ_chrY v1.3. A complete catalogue of the deposited files is provided in **Table S4**.

## Technical Validation

### Genome assembly quality assessment

To evaluate the base-level accuracy of the assembly, we utilized Merqury v1.3^32^ based on 21-mers derived from the PacBio HiFi reads (**Table 1**). The analysis yielded a QV of 57.54 (**Table 2**). This score notably exceeds the median QV of 48.0 reported for 43 diverse human Y chromosome assemblies^7^, demonstrating the exceptional sequence accuracy of our assembly. Assembly continuity, base-level accuracy, and structural integrity across the complete sequence were systematically evaluated. Genome-wide read coverage profiles confirmed uniform and continuous mapping of PacBio HiFi and ONT UL reads aligned using minimap2 v2.30 and Winnowmap v2.03 (**Figure S5**). CRAQ v1.10^33^ assessment report demonstrated exceptional quality metrics, including a 100% coverage rate, a low-confidence rate of 0, and an overall AQI of 99.781. Detailed inspection of a potential Clip-based Regional Error (CRE) identified at locus 58,323,159 (in the Yq12 region) revealed that while alignment via Winnowmap v 2.03 exhibited localized soft-clipping artifacts, realignment with VerityMap^34^ for both HiFi and ONT UL reads confirmed structural correctness and continuous alignment at this locus, validating CQ_chrY v1.3 as a high-fidelity assembly for downstream analyses.

### Terminal telomere integrity and absence of N-gaps

The structural completeness of the chromosomal termini was established by identifying canonical (TTAGGG)_n_ telomeric repeat arrays at both ends using seqtk telo (v1.4; https://github.com/lh3/seqtk). The 5’ PAR1 telomere spans 10,501 bp of continuous telomeric repeat units, whereas the 3’ PAR2 telomere spans 1,817 bp. Furthermore, in silico gap analysis using seqtk gap confirmed the complete absence of ambiguous *N* bases across the entire 68.50 Mb assembly.

### Data availability

The raw sequencing data have been deposited in the GSA-Human at the NGDC, CNCB, under accession number HRA017737. The CQ_chrY v1.3 genome assembly is available in the GenBase database at the same center, under accession C_AA501122.1. PacBio HiFi reads and Illumina short reads have been deposited in the GSA-Human at NGDC, CNCB, under accession number HRA001859^35^.

### Code availability

All custom scripts, command-line parameters, and configuration files used in this study are freely available in the GitHub repository at https://github.com/BoWangXJTU/CQ_chrY.

## Author contributions

X.Y., K.Y. and L.D. conceived this project; Y.Z., X.W. and L.D. collected samples; B.W., Z.Z., S.W. and P.Z. performed data analysis, curated the data, and wrote the manuscript. B.W. and X.Y. revised the manuscript. All authors have read and approved the final manuscript for publication.

## Ethical approval

The Quartet Project received approval from the Institutional Review Board of the School of Life Sciences at Fudan University (BE2050). The study was conducted in accordance with the principles outlined in the Declaration of Helsinki.

## Supplementary Material

**Figure S1. Assembly pipeline and dot plot validation of the draft CQ_chrY v1.2 genome.**

**Figure S2. Gap-filling strategy and breakpoint validation for the complete CQ_chrY v1.3 assembly.**

**Figure S3. Sequence synteny of PAR1, PAR2, and ampliconic regions in CQ_chrY v1.3 assembly.** PAR, pseudoautosomal region; AMPL, ampliconic; P1, Palindrome 1; P2, Palindrome 2; P3, Palindrome 3.

**Figure S4. Whole-chromosome sequence synteny between CQ_chrY v1.3 and other East Asian Y chromosome assemblies.**

**Figure S5. Whole-genome quality assessment and accuracy validation of the CQ_chrY v1.3 assembly.**

**Table S1. Hi-C sequencing statistics and contact-map quality metrics.**

**Table S2. Structural annotation of Yq12 heterochromatic repeat elements in the CQ_chrY v1.3 assembly.**

**Table S3. Annotation and sequence identity of genes on CQ_chrY v1.3 assembly.**

**Table S4. Complete catalogue of deposited raw data and assembled genome files.**

