## Supplementary figures and images for "Complete Telomere-to-Telomere Assembly of the Y Chromosome in the Chinese Quartet"

### Supplemental Figure 1

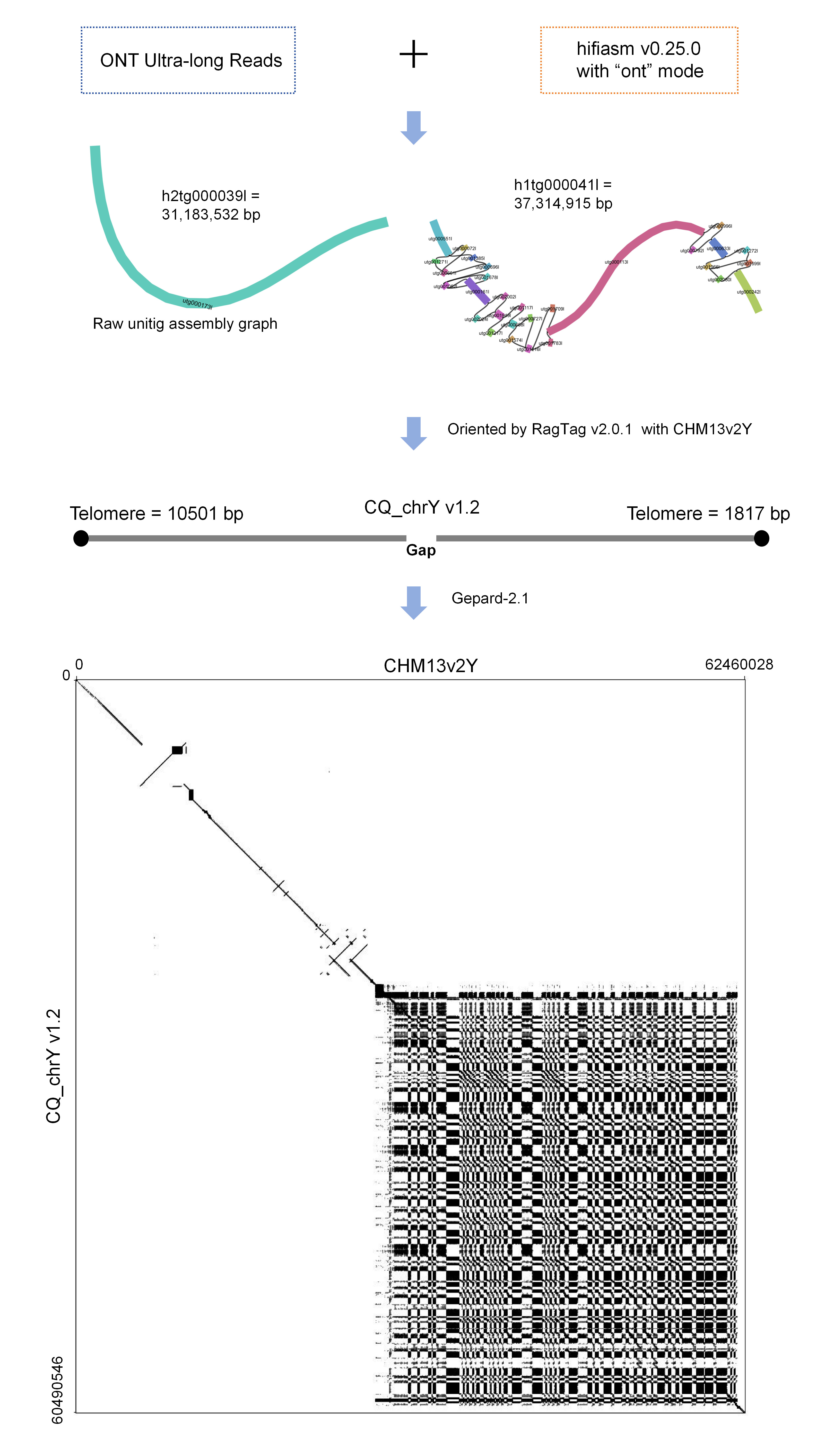

### Supplemental Figure 2

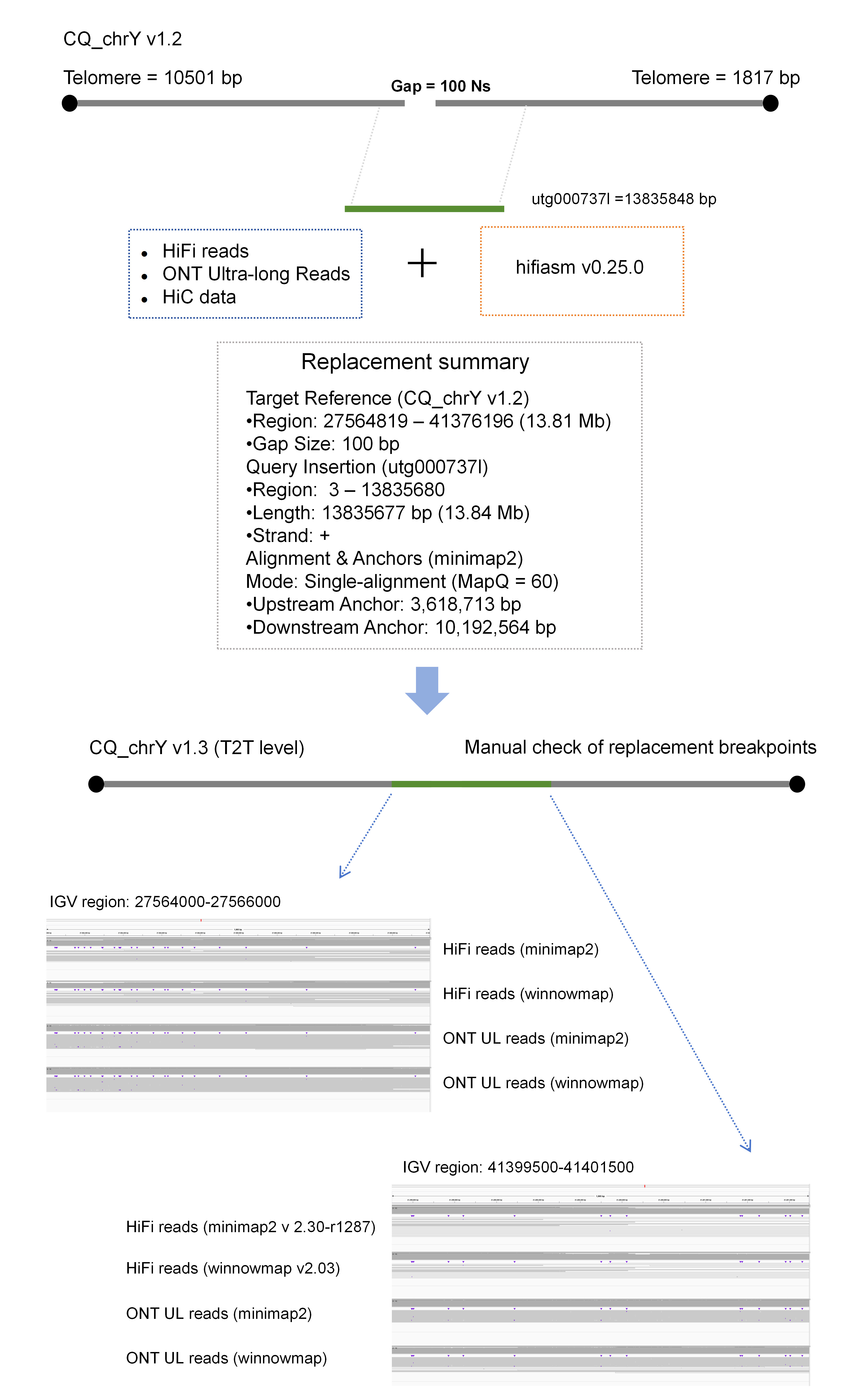

### Supplemental Figure 3

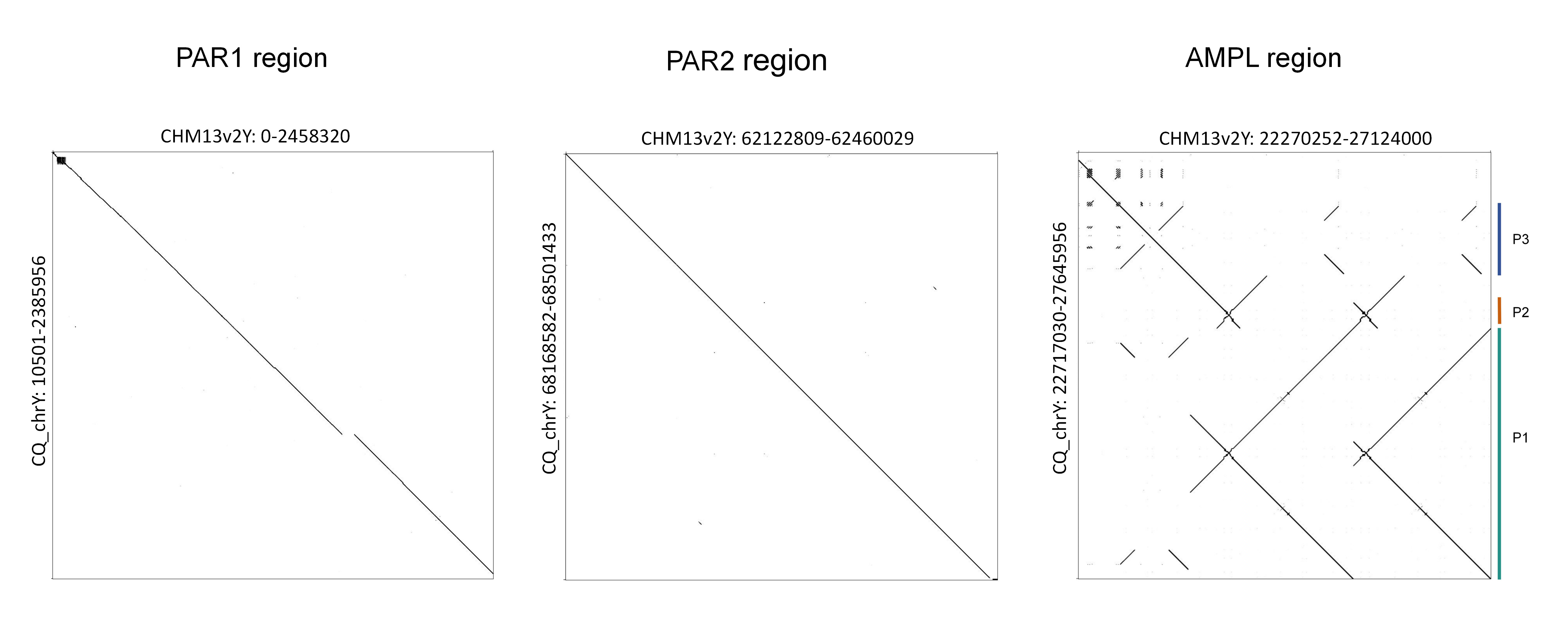

### Supplemental Figure 4

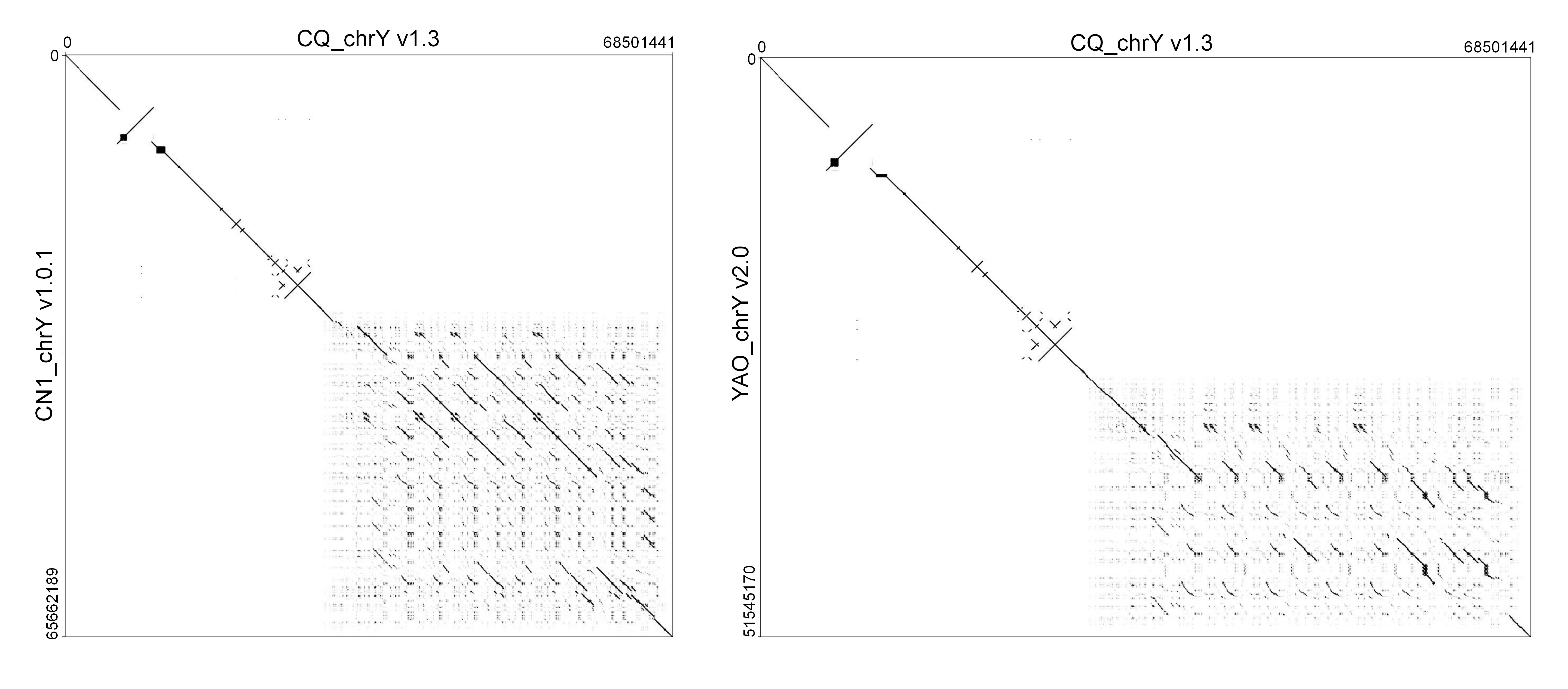

### Supplemental Figure 5

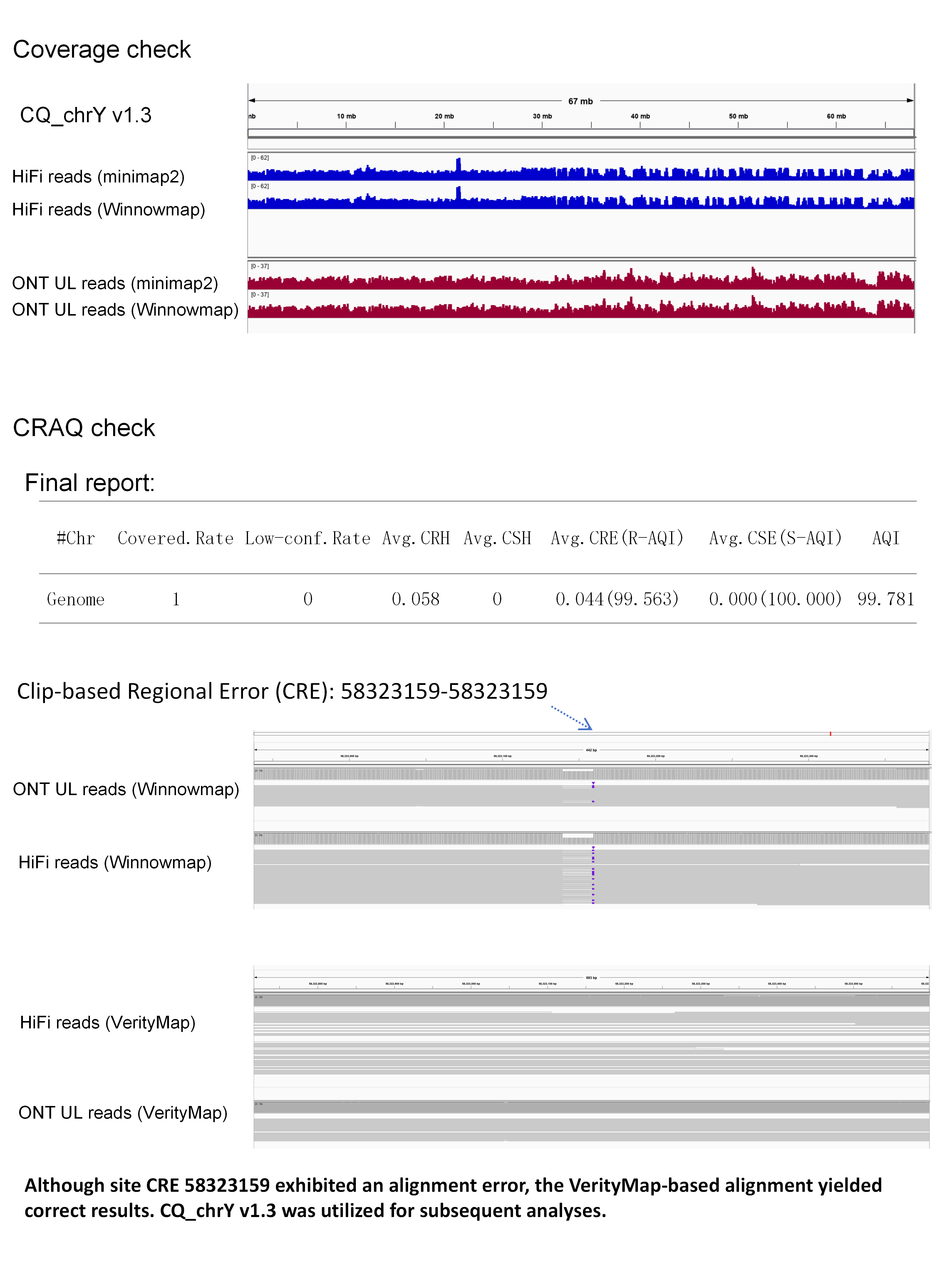
